# The impact of visual gaze direction on auditory object tracking

**DOI:** 10.1101/100065

**Authors:** Ulrich Pomper, Maria Chait

## Abstract

Subjective experience suggests that we are able to direct our auditory attention independent of our visual gaze, e.g when shadowing a nearby conversation at a cocktail party. But what are the consequences at the behavioural and neural level? While numerous studies have investigated both auditory attention and visual gaze independently, little is known about their interaction during selective listening. In the present EEG study, we manipulated visual gaze independently of auditory attention while participants detected targets presented from one of three loudspeakers. We observed increased response times when gaze was directed away from the locus of auditory attention. Further, we found an increase in occipital alpha-band power contralateral to the direction of gaze, indicative of a suppression of distracting input. Finally, this condition also led to stronger central theta-band power, which correlated with the observed effect in response times, indicative of differences in top-down processing. Our data suggest that a misalignment between gaze and auditory attention both reduce behavioural performance and modulate underlying neural processes. The involvement of central theta-band and occipital alpha-band effects are in line with compensatory neural mechanisms such as increased cognitive control and the suppression of task irrelevant inputs.

## Introduction

Humans can direct the locus of their auditory attention independently of visual gaze direction. While the two are often spatially aligned, there are many instances in everyday life in which we listen to somewhere else than where we are looking at. For instance, while driving and gazing toward the road ahead of us, we can engage in a conversation with someone sitting next to or behind us. Here, we seek to understand how gazing toward versus away from the locus of auditory attention affects behavioural and neural responses to sounds, and its impact on more global measures of brain states such as ongoing oscillatory activity.

Visual gaze is usually an overt manifestation of selective visual attention^1,2^, and often tightly linked with attention in other modalities^3,4^. During dichotic listening tasks, spontaneous eye movements have been shown to occur preferentially toward the attended side^5,6^. Research in animals has shown that the direction of visual gaze modifies concurrent auditory processing. For example, Werner-Reiss et al.^7^ found that eye position changes both the spontaneous activity and responses to sounds of neurons in the auditory cortex of awake macaques, even in complete darkness. Similarly, Groh et al.^8^ demonstrated that in macaques eye position affects firing rates of auditory neurons already at the level of the inferior colliculus. In humans, research on the impact of gaze direction on auditory processing has mostly been limited to its effects on sound localization. For instance, Maddox et al.^9^ reported that directing gaze toward a sound significantly enhances discrimination of both interaural level and time differences, whereas directing auditory spatial attention alone does not.

Irrespective of gaze direction, a large number of electroencephalographic (EEG) studies in humans have shown that endogenous auditory attention can amplify event related potentials (ERPs) to sounds as early as 20 ms after stimulus onset^10–13^. These early attentional effects are thought to reflect a sensory selection mechanism, based on readily discriminable features such as spatial location^10,14^. In addition to unisensory auditory attention, covert visual attention to the location of a sound can both amplify ERPs, as well as facilitate behavioural responses to auditory targets^15–17^, demonstrating the potential impact of visual information on auditory processing. Apart from influencing phasic responses to single sounds, auditory attention also affects oscillatory neural activity in the alpha-band range (8-14 Hz)^18,19^. For example, Obleser and Weisz^19^ presented human listeners with degraded speech, and found that occipital alpha band activity correlated with speech intelligibility and listening effort. More generally, alpha-band activity has been suggested to act as a local sensory gating mechanism, by which processing of relevant sensory inputs is enhanced and irrelevant input is suppressed^20^. Indicative of this, both visual and auditory spatial attention have been shown to induce lateralized changes in occipital alpha-band activity, with larger power contra-versus ipsilateral to the attended side^21–23^. While the topography of alpha-band modulation for visual and auditory attention overlaps, the underlying networks for the two modalities are likely distinct^18,21^. Prolonged attention and demanding cognitive performance has additionally been linked to increased central theta-band activity (4-7 Hz)^24,25^, and has been demonstrated for tasks in the auditory^26^, and visual^27^ domain, as well as during multisensory processing^28^. For instance, Friese et al.^28^ found increased fronto-medial theta-band activity during attended versus unattended trials in an audio-visual congruency detection paradigm. In summary, auditory attention in humans increases ERPs to sounds and modulates posterior alpha-band oscillations, while the amount of cognitive control required during a given task seems to be reflected in central theta-band activity.

However, while there is increasing knowledge about the mechanisms of auditory attention as well as the mediating influence of visual gaze in other animals, the impact of gaze direction on human auditory processing is still largely unknown. To our knowledge, the only previous study investigating gaze dependent changes in the quality of auditory processing in humans was performed by Okita and Wei^29^ Using EEG recordings from four electrodes, the authors show enhanced ERPs between 100 and 500 ms following a tone, when participants gazed towards versus away from the spatial source of the tone. This was interpreted as an increase of selectivity between relevant and irrelevant auditory inputs. However, this experiment suffers from a number of methodological limitations including the lack of quantifiable eye position monitoring. Furthermore it did not find any behavioural effects of gaze direction on auditory processing.

In the present EEG study, we used a full factorial design to investigate the impact of task irrelevant gaze direction and attention, as well as their interaction, on auditory processing. Participants attended to target sounds presented from one of three loudspeakers, while either gazing at the same or a different loudspeaker. We hypothesized that gazing toward to location of auditory attention (*coherent* condition) will lead to improved behavioural performance, as well as changes in ERP responses and power of neural oscillations, compared to when gazing away from the location of auditory attention (*incoherent* condition).

## Methods

### Participants

Nineteen paid volunteers (10 females, mean age 25.3 years) participated in this study. All were right handed (Edinburgh Handedness Inventory^30^) and reported no neurological illness or hearing deficits. An additional participant was excluded from analysis due to extensive muscle and eye-movement artifacts. The study was approved by, and conducted in accordance with the research ethics committee of the University College London, and all participants provided written informed consent.

### Task and Procedure

Participants were seated in an acoustically shielded room (IAC Acoustics, Hampshire, UK), with their head fixed in a headrest. Three loudspeakers were placed 80 cm away from the participants’ head, and vertically located at the level of the ears. The left loudspeaker was located at −30 degrees, the central loudspeaker at 0 degrees and the right loudspeaker at −30 degrees of the participants head (see Figure 1A). The main experiment consisted of 32 trials with a duration of one minute each. Prior to each trial, participants received instructions regarding which loudspeaker to attend and which loudspeaker to gaze at via an LCD screen, which was located behind the loudspeakers. These instructions remained on the screen throughout the trial. Additionally, a red LED, attached at the centre of each loudspeaker, indicated which loudspeaker to gaze at. Participants were instructed to fixate the LED throughout the duration of each trial.

**Figure 1:**
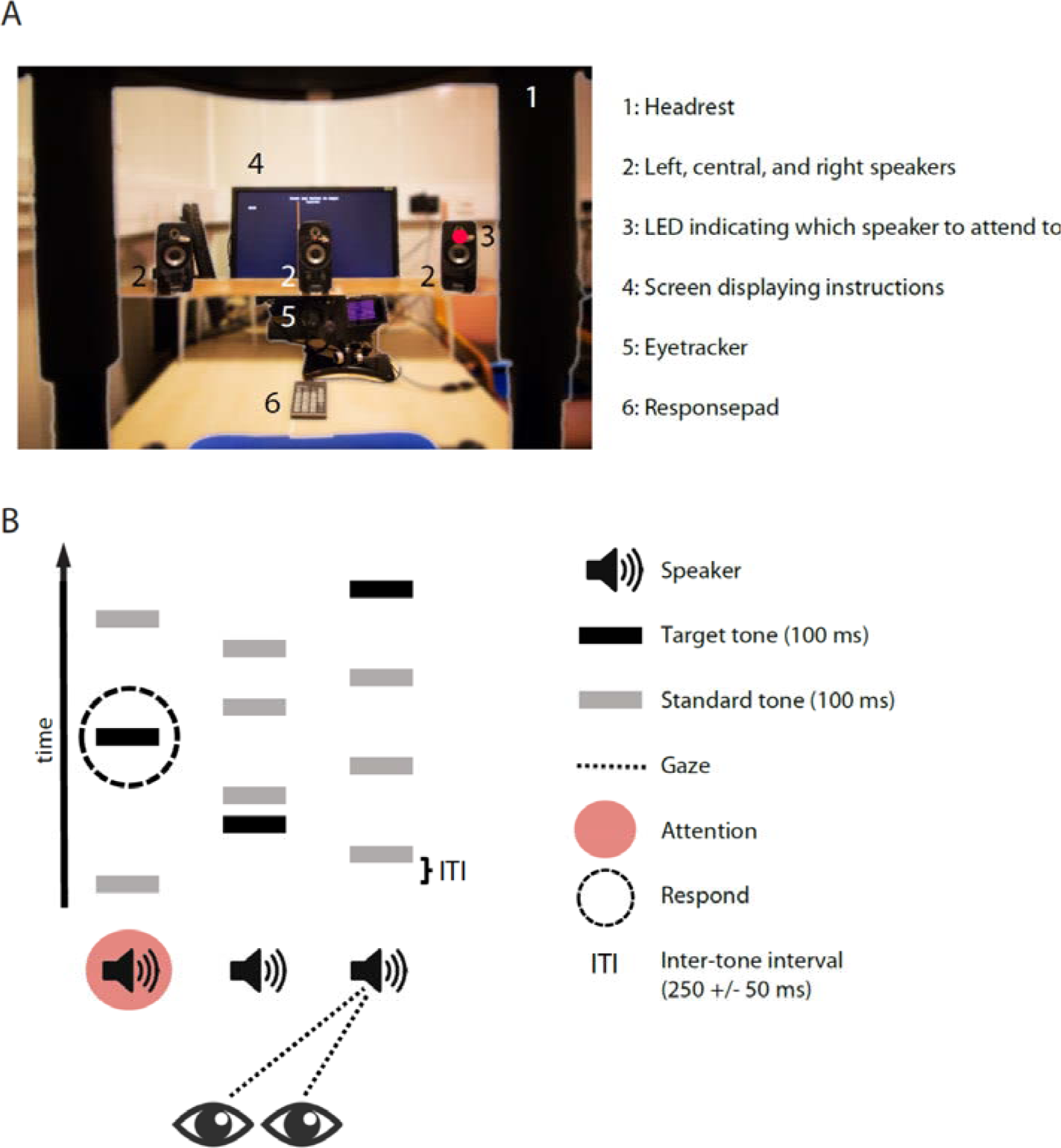
Setup and trial design. (A) View from the participants’ perspective. (B) Schematic experimental design. Continuous streams of sounds were simultaneously presented from each of the three loudspeakers. The sound streams consisted of both standard and target auditory stimuli. On each trial, participants were instructed to attend one of the three loudspeakers (here: left speaker), and visually gaze at either the same of a different speaker (here: right speaker). The task was to provide speeded responses to target sounds coming from the attended location.

During each trial, streams of pure tones were presented from the three loudspeakers (left, right and central; Figure 1B). Each tone had a duration of 100 ms (5 ms rise and fall) and a frequency of 660 Hz. The interstimulus interval (ISI) between successive tones (independent of loudspeaker location) was jittered between 200 to 300 ms (mean 250 ms). The location (loudspeaker) the tones were presented from was randomized, with the restriction that no more than 3 successive tones could be presented from the same loudspeaker. Effectively, the stimulus was perceived as three concurrent streams, each with a random ISI. Twenty percent of all tones were amplitude modulated at 20 Hz, and these were designated as target tones. The participants’ task was to provide a speeded response with their right index finger, to target tones presented from the attended loudspeaker only. Target tones presented from the two unattended loudspeakers had to be ignored. In total, each trial contained 44 tones from each loudspeaker (132 altogether), 8 of which were targets (24 altogether). The combination of three possible gaze directions and three possible attentional locations resulted in nine different experimental conditions. For each condition, a block of four consecutive trials was presented. The order of blocks was randomized across participants.

### Eyetracking

To ensure correct fixation throughout the experiment, the direction of gaze was continuously monitored using an eye tracker (Eyelink 1000, SR Research Ltd., Ontario, Canada). Data were recorded binocularly with a sampling rate of 500 Hz. Prior to each experimental block, the system was calibrated using a 9 point calibration procedure. Offline, eyeblinks as well as periods of missing data were removed from the analysis^31^. Figure 2 shows a descriptive heatmap of eye position locations, with data pooled across all participants and conditions. To better illustrate fixations, the data are plotted on top of an image of the visual field containing the three loudspeakers. Additionally, the figure contains histograms of all the fixation locations along the horizontal and vertical axis. As can be seen, fixations were centred around the instructed location on each loudspeaker. Note that small inconsistencies existed between subjects in the calibration and the exact placement of the fixation-indicating LED, which led to an increased variance of the data in the heatmap and the corresponding histograms. As a more accurate measure of the within-subject variability in fixation-location, we calculated the full-width at half maximum (FWHM) of the fixation histograms, separately for each participant and condition. The average FWHM was 18.5 pixels, with a range from 7.3 to 35.5 pixels between participants and conditions. This was well within the spatial extend of the face of each loudspeaker, which was corresponding to 173 x 702 pixels. To rule out potential differences in average horizontal fixation location as well as fixation FWHM between conditions, we conducted a 3-way ANOVA using the factors Attention location (left, centre, right), Gaze location (left, centre, right) and eye (left, right). For fixation location, we found an obvious and expected main effect of Gaze location (F(1,36) = 8398, p < .001). Importantly, we observed no additional differences between conditions (all p > 0.13). For FWHM values, no main effects or interactions were apparent (all p > 0.12). Thus, we can assume that participants complied with the gaze instructions, and that no systematic differences in fixation behavior between conditions are present.

**Figure 2:**
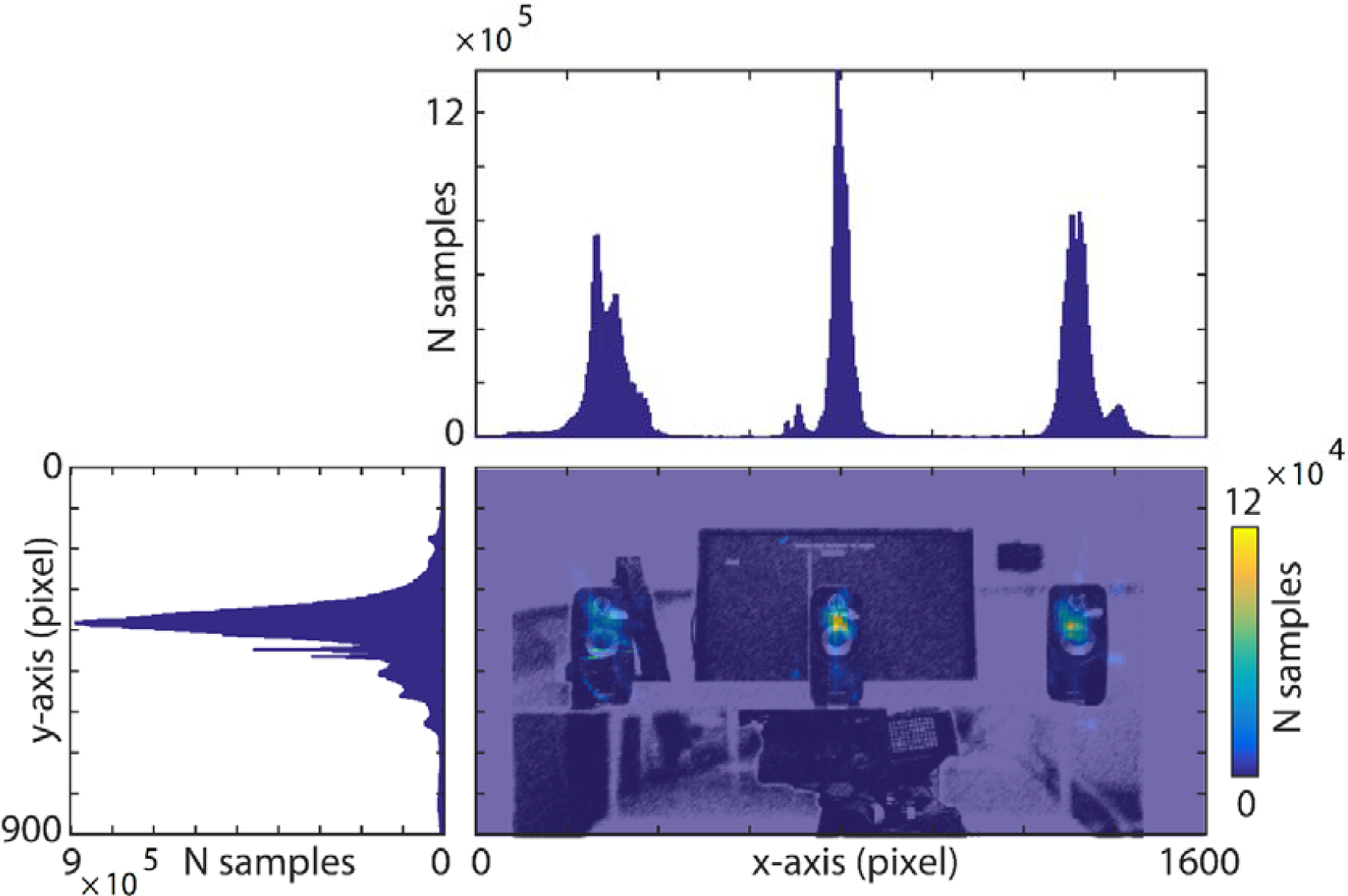
Eye-fixation data. Central plot: Heatmap of visual fixation duration, overlaid on the part of the visual field which contained the three loudspeakers. Heatmap and picture are corresponding in size. The data shown are pooled across all subjects and conditions. Warm colours indicate a large number of samples during which a point was fixated. Upper plot: Histogram showing the number of samples each point in the horizontal axis was fixated. Clear peaks for the left, central and right loudspeaker location are visible. Left plot: Histogram showing fixations for the vertical axis.

### EEG recording and data preprocessing

High-density EEG was recorded from 128 scalp channels, using a Biosemi ActiveTwo system. To monitor eye movements, two additional electrodes were placed at the medial upper and lateral border of the right ocular orbit. Recordings were made reference-free with a passband of 0.016–250 Hz and digitized at a sampling rate of 2048 Hz. All off-line data processing was done using EEGLAB (http://www.sccn.ucsd.edu/eeglab^32^) and FieldTrip (http://www.ru.nl/fcdonders/fieldtrip^33^), implemented in Matlab (The MathWorks Inc., Natick, MA, USA). Off-line, data were bandpass filtered (using a finite impulse response filter) between 0.3 and 125 Hz, downsampled to 256 Hz and re-referenced to common average. An additional narrow-band notch filter (49.5–50.5 Hz, 4th order zero-phase Butterworth filter) was applied to remove remaining line noise. Trials containing muscle- and technical artifacts were removed by visual inspection. On average, less than 1 % of trials were removed. Electrodes with extremely high- and/ or low-frequency artifacts throughout the recording (mean = 1.1) were linearly interpolated using a model of the amplitude topography at the unit sphere surface based on all nonartifactual electrodes^34^. To reduce artifacts such as eye-blinks, horizontal eye movements, and electrocardiographic activity, an independent component analysis approach was applied (extended Runica^35^). Components representing artifacts were removed from the EEG data by back-projecting all but these components (mean = 6.1). Finally, continuous data were cut into epochs from −100 ms to 600 ms around each tone onset, and baseline-corrected by subtracting the mean pre-stimulus activity between −100 to 0 ms.

### Statistical analysis of behavioral data

Dependent measure were d’ sensitivity scores^36,37^ and reaction time (RT) computed from target tone onset. Prior to the analysis, for each subject, RTs above or below 3 standard deviations of the condition mean were excluded from the analysis. We also analyze the false alarm rate (FA) as a measure of distractibility, as gazing away from the attended location might particularly increase responses to task irrelevant target tones coming from the gazed-at location. RTs, d-prime values and FAs were compared between the experimental conditions using 2-way repeated measures ANOVAs with the factors Spatial Coherence (*coherent* vs. *incoherent*) and Loudspeaker Location (*left* vs. *central* vs. *right*).

### Analysis of Event-related potentials

For the analysis of ERPs to individual sounds, we were interested in the main effect of attention, the main effect of gaze, as well as a potential interaction between the two factors. To rule out confounding motor artifacts associated with target trials, our main ERP analysis was performed on epochs containing standard sounds only. However, we performed the same analysis procedure separately for epochs containing target trials as well. As a first step, we defined analysis time-windows based on a grand-average ERP, pooling across all conditions (Figure 3A). The grand average ERP revealed a standard response consisting of P1 (65 ms), N1 (125 ms), P2 (175 ms) peaks^12,38,39^. We defined the time-windows as: 50 to 80 ms (P1), 110 to 140 ms (N1), and 160 to 190 ms, (P2) (i.e. +/- 15 ms around the local peak of the potential). We then proceeded with the analysis using two independent complementary approaches. In the first, more data-driven approach, we computed cluster-based permutation tests within each of the three time windows, separately for the main effect of Attention (attended vs. unattended), the main effect of Gaze (gazed-at vs. not gazed-at), and their interaction (attended and gazed-at minus attended and not gazed-at vs. unattended and gazed-at minus unattended and not gazed-at). The cluster-based permutation-tests comprised of pairwise t-tests between the conditions, conducted for each time-point (within the predefined time-windows) and channel. This procedure controls the type I error rate in statistical tests involving multiple comparisons by clustering adjacent data points exhibiting the same effect^40^. The threshold of the dependent samples t-tests and the permutation P-value of the cluster were both set to p = 0.05, and 1000 permutations were calculated for each comparison. For the second approach, we selected a fronto-central region of interest (ROI) based on the topography of our present P1, N1 and P2 peaks (Figure 3A), and in line with numerous previous studies^38,39,41,42^. Note that this ROI consisted of many channels which were also present in the channel clusters independently obtained with the first, data driven analysis approach. We then averaged ERP amplitudes within this ROI separately for each condition, time-window, and participant. Finally, we conducted a repeated measures ANOVA for each of the three time-windows (P1, N1, P2) using the factors Attention (attended vs. unattended) and Gaze (gazed-at vs. not gazed-at).

**Figure 3:**
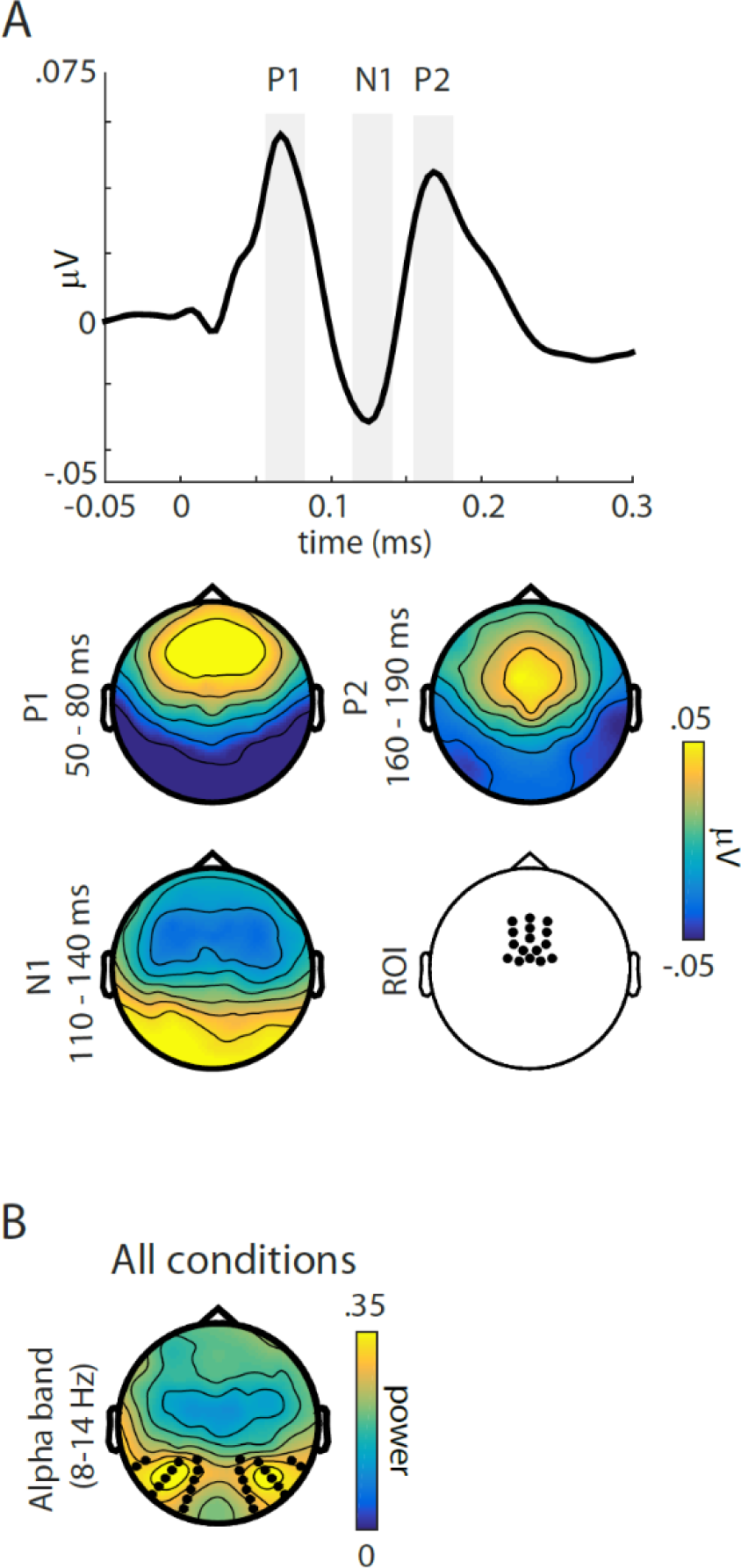
Definition of regions of interest (ROIs) for subsequent analyses. **(A)** Left side: Grand average ERP trace across all conditions, using a fronto-central ROI. Grey bars indicate time-windows of interest, selected around (+/- 15 ms) the prominent P50, N1 and P2 peaks. Right side: Grand average topographies across all conditions of the three selected time-windows of interest, as well as topography indicating the pre-selected fronto-central ROI. **(B)** Topography of grand-average alpha-band power (8-14 Hz) across all conditions. Black dots indicate the channels used for statistical analysis.

Although differences between loudspeaker locations were not our main focus of analysis, we also ran an additional 3 way ANOVA with the extra factor of Loudspeaker Location (left, centre, right), to investigate potential differences between the spatial origin of sounds. Further, previous studies on sound localization have shown that the perceived sound location is steadily shifted toward the direction of gaze over longer periods of fixation^43^. To investigate potential dynamic changes in the effect of gaze onto ERPs to sounds, we divided the data of each condition into chunks of 20 seconds. Since each trial lasted one minute and four trials of the same condition were presented consecutively, this resulted in 12 consecutive analysis periods. We compared these time-periods using a 3-way ANOVA with the factors Attention (attended vs. unattended) and Gaze (gazed-at vs. not gazed-at) and Time (periods 1 to 12).

### Analysis of oscillatory brain activity

Here, we were interested in the overall differences between conditions in which attention and gaze were spatially aligned (coherent) versus conditions where attention and gaze were spatially misaligned (incoherent). Particularly, we compared the ‘attend-left, gaze-left’ to the ‘attend-left, gaze-right’ condition, and the ‘attend-right, gaze-right’ to the ‘attend-right, gaze-left’ condition. The ‘attend-central, gaze-central’ condition was compared to both the ‘attend-central, gaze-left’ condition (referred-to as central^L^) and to the ‘attend-central, gaze-right’ condition (referred-to as central^R^) separately. As a first step, we transformed activity within individual epochs into the frequency domain by applying a Fast Fourier Transform with a single Hanning taper. Power at frequencies from 2 to 30 Hz was computed in 0.5 Hz steps, using a fixed frequency smoothing (f = 2 Hz). In line with previous work on auditory and spatial attention^18,44,45^ the main focus of our analysis was activity in the alpha-band frequency range (8-14 Hz). Next, we defined a ROI which best reflect both attention- and gaze related changes in alpha-band activity. Since, for the present study, differences in alpha-band power between conditions were strongly lateralized to the left or right side (see Figure 7B), defining a ROI based on the average difference between conditions was not feasible. Thus, we defined ROIs by averaging alpha-band activity across all conditions, and selecting 26 occipital channels (13 on each side), which exhibited the most robust alpha-band activity (Figure 3B). Importantly, the ROIs defined by this procedure overlap nicely with the topography of the attention- and gaze dependent alpha-band modulations shown in Figure 7 A. Alpha-power within these ROIs was then converted into the alpha-modulation index 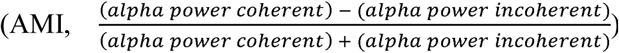, which represents a normalized difference of alpha power between the coherent versus incoherent trials^46,47^. This normalization step is crucial, since we compare absolute spectral power taken from different blocks of the experiment. Finally, AMI was averaged across all channels within the left and right ROI, and entered into a 2-way repeated measures ANOVA using the factors Attention Location (*left* vs. *centra^R^* vs. *central^L^* vs. *right*) and ROI (*left* vs. *right*). Significant 2-way interactions were further investigated by comparing AMI between left and right ROIs separately for each Attention Location, using t-tests. Further, we calculated one sample t-tests for each condition and location to test whether the respective AMI differs significantly from zero.

In addition to alpha-band activity, we were also interested in overall differences in theta-band (4-7 Hz) power between the coherent and incoherent conditions. Similar to the alpha-band analysis, we set out to define a spatial ROI reflecting attention- and gaze related changes in theta-band activity. However, unlike alpha-band activity, the topographical distribution of theta-band activity was not lateralized but highly similar between conditions (irrespective of the spatial coherence or the attentional location, see Figure 8A). Due to this fact, we based the theta-band ROI on the contrast between all coherent versus all incoherent conditions, pooled together across the three attention locations (left, center, right). This comparison was done by means of a cluster-corrected permutation test, with 1000 permutations and the threshold of the dependent samples t-tests and the permutation p-value of the cluster set to p = 0.05^40^. Figure 8A shows the ROI defined by this procedure, comprising of 36 channels and located over central and posterior parts of the scalp. Analogous to the alpha-band and ERP analysis, we then conducted a second level analysis to investigate the effect of coherence separately for attending to the left, central, and right loudspeakers. Theta-band power was *(theta power coherent) - (theta power* converted into theta-modulation index 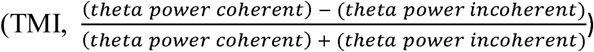 averaged across the 36 channels/ ROI separately for each condition and participant, and entered into a repeated measures ANOVA with the factor Attention Location (*left* vs. *centraf* vs. *central*^L^ vs. *right*). Additional one sample t-tests were calculated for each condition to test whether TMI differs significantly from zero.

### Correlation analysis

As an exploratory measure, we were interested in a potential correlation between behavioural data and EEG responses. In order to reduce the number of statistical comparisons, we first computed an index of effect of spatial coherence by calculating the difference values between coherent and incoherent conditions for RTs as well as a modulation index for spectral power 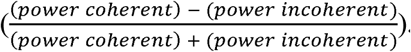. We then calculated pointwise Pearson s correlations between these measures in RTs and spectral power, at all channels and frequencies between 2-30 Hz.

## Results

### Behavioural results

Figure 4 shows RTs, d-prime values and FA rates for the different experimental conditions. The 2-way ANOVA for RTs using the factors Spatial Coherence (*coherent* vs. *incoherent*) and Locations (*left* vs. *center* vs. *right*) yielded a significant main effect of Spatial Coherence (F_(1,18)_ = 7.11, p = .016). Participants were overall faster (mean = 17 ms) when responding during the *coherent* compared to the *incoherent* condition. No main effect for location and no interaction was found (p values >.092). For FA rates, the ANOVA yielded a significant main effect of Spatial Coherence (F_(1,18)_ = 5.85, p = .026) as well as of Location (F_(1,18)_ = 6.96, p = .003), but no interaction (p>.051). Interestingly, participants had a higher FA rate during the coherent compared to the coherent condition (see discussion below). No significant effects were found for d-prime values (all p values >.067).

**Figure 4:**
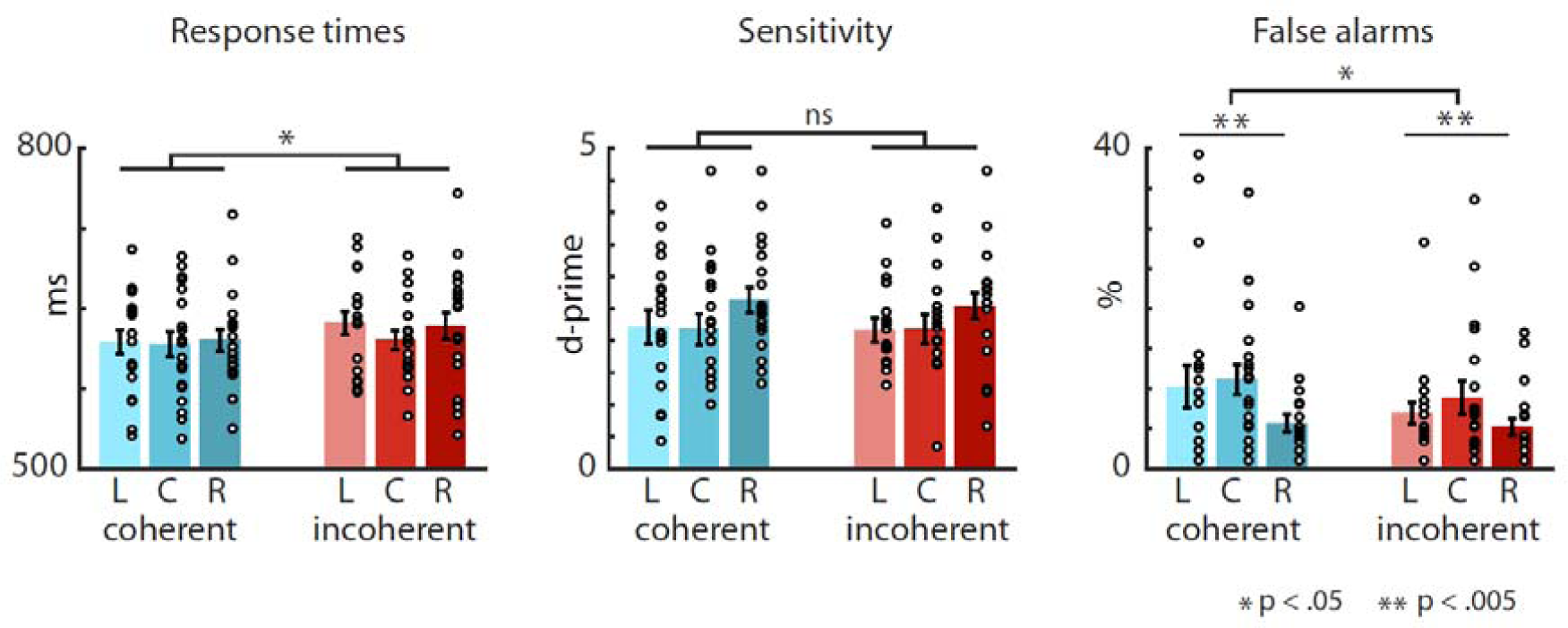
Behavioural results. Mean response times (left barplot), d-prime values (middle barplot), and flase-alarm rates (right barplot) to targets presented from the left (L), center (C) and right (R). Blue hues indicate coherent conditions (attention and gaze towards the same location), red hues indicate incoherent conditions (attention and gaze towards different locations). The dots represent the individual participants’ performance.

### Event-related potentials to standards

Figure 5 shows the results for the analysis of ERP to standard sounds, in which we investigated main effects of Attention, Gaze, and their interaction. Figure 5A displays ERP traces collapsed across the left, central and right loudspeaker location. For each condition, a prominent P1 (~ 65 ms poststimulus), N1 (~ 125 ms poststimulus), and P2 (~ 175 ms poststimulus) peak is present. The overall shape and peak latencies are similar for each of the conditions. Attended compared to unattended sounds show larger P1 and N1 peak amplitudes, as well as lower P2 amplitude. Interestingly, no differences in amplitudes are seen for the gaze versus no-gaze comparison.

**Figure 5:**
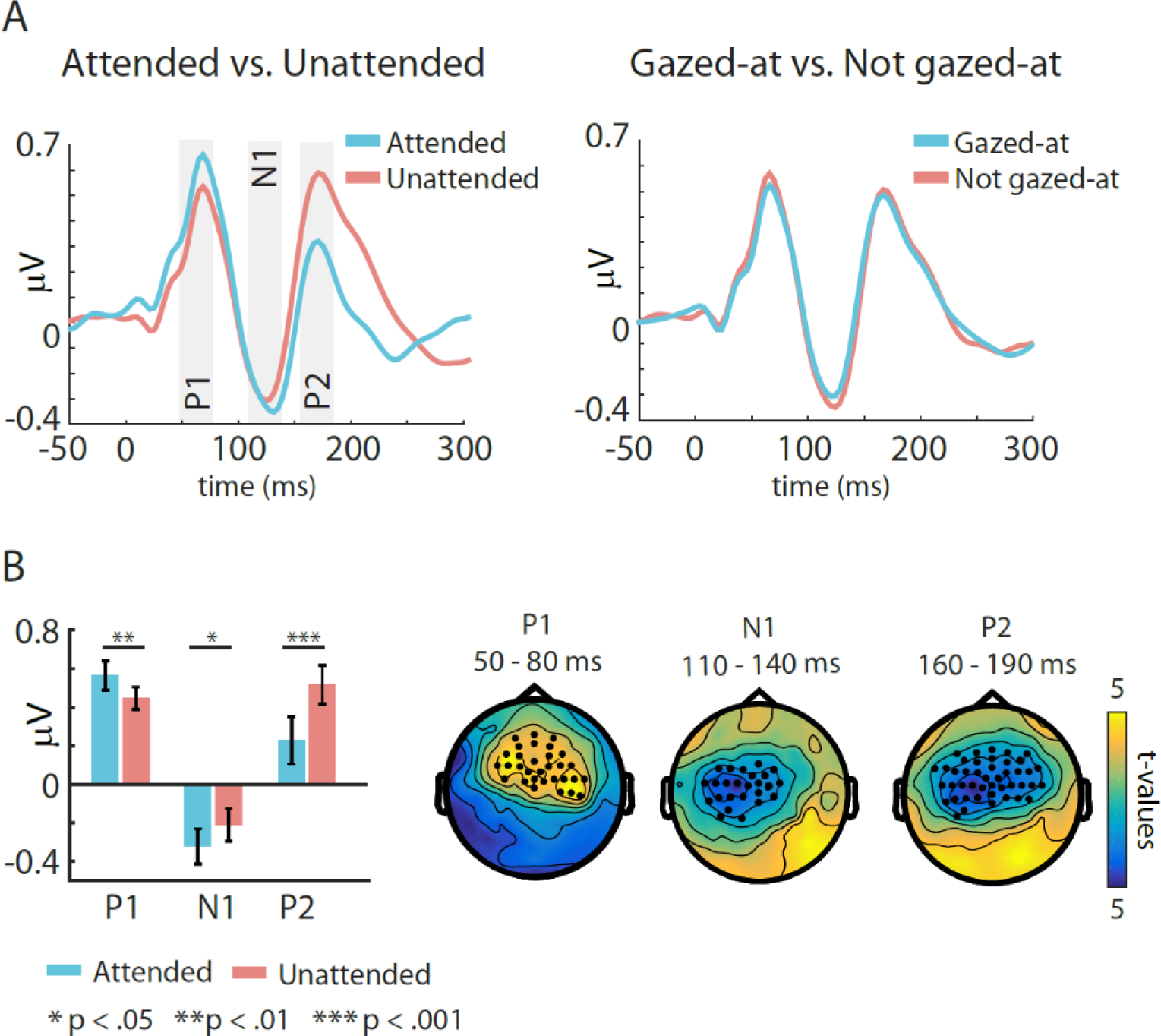
Grand average event-related potentials to standards. **(A)** Grand average ERP results using a predefined fronto-central ROI (Figure 3A). Shown are traces for the Attended (blue colour) versus Unattended (red colour) conditions (left), as well as the Gazed-at (blue colour) versus Not gazed-at (red colour) conditions (right). Significant results for the investigated three time-windows (indicated by grey bars) were found only for the Attended versus Unattended comparison. **(B)** Statistical results for the Attended versus Unattended comparison. Left: Barplot showing the results for the ANOVA using a predefined ROI (Figure 3A). Right: Topographies showing the results of the cluster-based permutation tests. Significant clusters of channels were found in all three time windows. Colours indicate t-values, black dots indicate channels belonging to the significant cluster.

In our first analysis approach, we found significant clusters of electrodes for the comparison between attended and unattended sounds for all three time-windows (Figure 5B, right topoplots). For the P1 and N1, the cluster was a result of larger amplitudes in the attended compared to the unattended condition (mean cluster t = 2.60, p < 0.019 and mean cluster t = - 3.05, p < 0.007, respectively). For the P2, the cluster was due to larger amplitudes in the unattended compared to the attended condition (mean cluster t = −4.54, p < 0.007). The topography of the clusters was overlapping, but slightly more anterior for the P1 compared to the N1 and P2. Importantly, we found no significant differences between the gazed-at and not gazed-at conditions, as well as no interactions between Attention and Gaze.

For our second analysis approach, we calculated a 2-way ANOVA with the factors Attention (attended vs. unattended) and Gaze (gazed-at vs. not gazed-at) for each of the three time-windows, using a predefined fronto-central ROI (Figure 5B, left barplot). For the P1 and N1 time windows, we found a significant main effect of Attention (F(1,18) = 8.58, p < .009 and F(1,18) = 7.83, p < .012, respectively), due to larger amplitudes in the attended compared to the unattended condition. For the P2 time window, we found a significant main effect of Attention (F(1,18) = 24.32, p < .001), due to larger amplitudes in the unattended compared to the attended condition (Figure 4A, left barplot). In line with the results from the first analysis approach, we found no significant main effects of Gaze (all p > .116) and no interactions (all p > .063).

When extending our two-factorial ANOVA by the factor Location (left vs. central vs. right), we found an additional main effect of Location for the N1 (F(1,36) = 5.31, p < .010), due to larger amplitudes at the right compared to the central and left loudspeakers (t(18) = −2.34, p < .035 and t(18) = −3.05, p < .067). We found the same main effects of Attention for the P1 (F(1,18) = 8.10, p < .011), the N1 (F(1,18) = 7.84, p < .012) and the P2 (F(1,18) = 24.93, p < .001) as in the main analysis. To investigate differences between loudspeaker location, we ran an extended 3-way ANOVA with the added factor Location (left vs. central vs. right). In addition to the same results found in the 2-way ANOVA, we also found a main effect of Location for the N1 (F(1,36) = 5.31, p < .010), due to larger amplitudes at the right compared to the central and left loudspeakers (t(18) = −2.34, p < .035 and t(18) = −3.05, p < .067). No other effects of Location were found. Finally, we investigated potential temporal dynamics in the effect of gaze on ERPs, by computing a three-way ANOVA using the factors Attention (attended vs. unattended), Gaze (gazed-at vs. not gazed-at) and Time (time period 1 to 12). In addition to the same main effects as in the 2-way ANOVA, we also found main effects of Time for the N1 and the P2 (F(1,18) = 7.4, p < .001 and F(1,18) = 4.31, p < .001, respectively), due to overall larger amplitude levels in the last compared to the first time period. Importantly however, we found no interactions between the factors Time and Attention or Time and Gaze (all p > .243), suggesting that passage of time had no impact on the effect of attention or gaze on ERPs.

### Event-related potentials to targets

Figure 6 shows the results for the analysis of ERPs to target sounds. The ERP traces (Figure 6A; collapsed across the left, central and right loudspeaker location) look similar to those from the standard sounds, with a more prominent increase of the P2 for the unattended targets. Larger P2 amplitudes are commonly observed for infrequent or target stimuli, and are larger for irrelevant compared to relevant stimuli^48,49^. Further, an early but not significant difference around the P1 is seen between the gazed-at and not gazed-at conditions. For the statistical analysis using the data driven cluster-based permutation approach, we found a significant cluster only for the attended versus unattended comparison during the P2 time-window (mean cluster t = −3.37, p < 0.007). Similarly, in the alternative approach using a predefined ROI, the two-way ANOVA with the factors Attention (attended vs. unattended) and Gaze (gazed-at vs. not gazed-at) yielded a significant main effect of Attention (F(1,18) = 19.45, p < .001) for the P2, due to larger amplitudes in the unattended condition. None of the other main effects or interactions were significant (all p > .21) Thus, although ERPs to targets showed similar topographies and traces as those to standard sounds, only one of the three effects found for standards was present in the target analysis. Given the low number of target sounds, we did not further separate the data to investigate effects of speaker location or passage of time.

**Figure 6:**
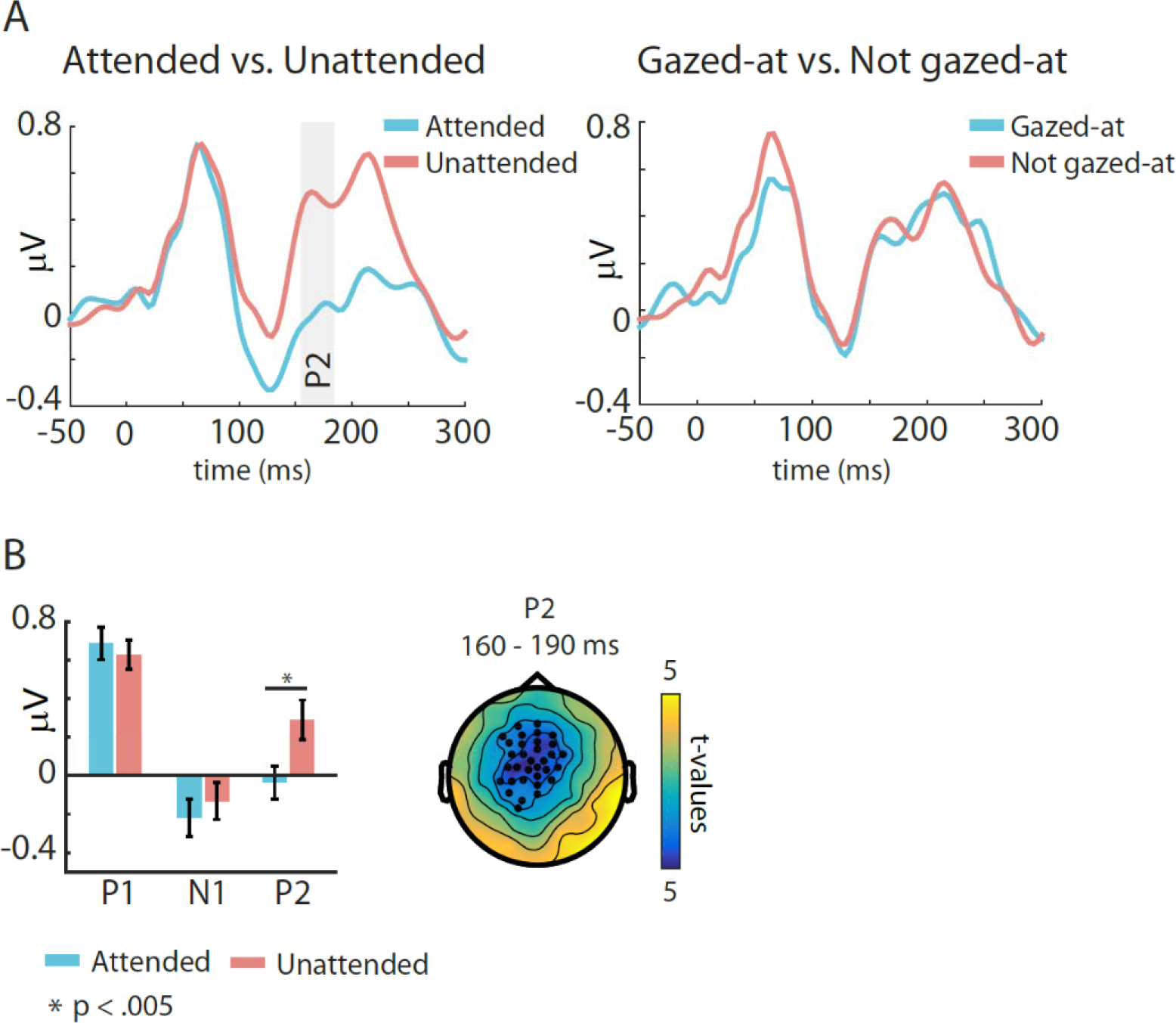
Grand average event-related potentials to targets. **(A)** Grand average ERP results using a predefined fronto-central ROI (Figure 3A). Shown are traces for the Attended (blue colour) versus Unattended (red colour) conditions (left), as well as the Gazed-at (blue colour) versus Not gazed-at (red colour) conditions (right). Significant results for the investigated three time-windows were found only for the P2 in the Attended versus Unattended comparison. **(B)** Statistical results for the Attended versus Unattended comparison. Left: Barplot showing the results for the ANOVA using a predefined ROI (Figure 3A). Right: Topography showing the results of the cluster-based permutation test. Significant clusters of channels were found for the P2 time window. Colours indicate t-values, black dots indicate channels belonging to the significant cluster.

### Oscillatory EEG activity

Figure 7A illustrates the scalp topography of the grand-average alpha-band activity (8-14 Hz) within each condition. Overall, alpha-band activity was most prominent at posterior sites, and appears to be stronger in the incoherent compared to the coherent condition for each of the four illustrated comparisons. Moreover, alpha-band power showed a pattern of lateralization depending on the location of both attention and gaze. Specifically, when attention and gaze are directed toward different sides (incoherent condition), alpha-band activity is increased at the hemisphere contralateral to the direction of gaze (Figure 7A, right column). This effect is particularly evident in topography of the AMI (Figure 7B, left column).

**Figure 7:**
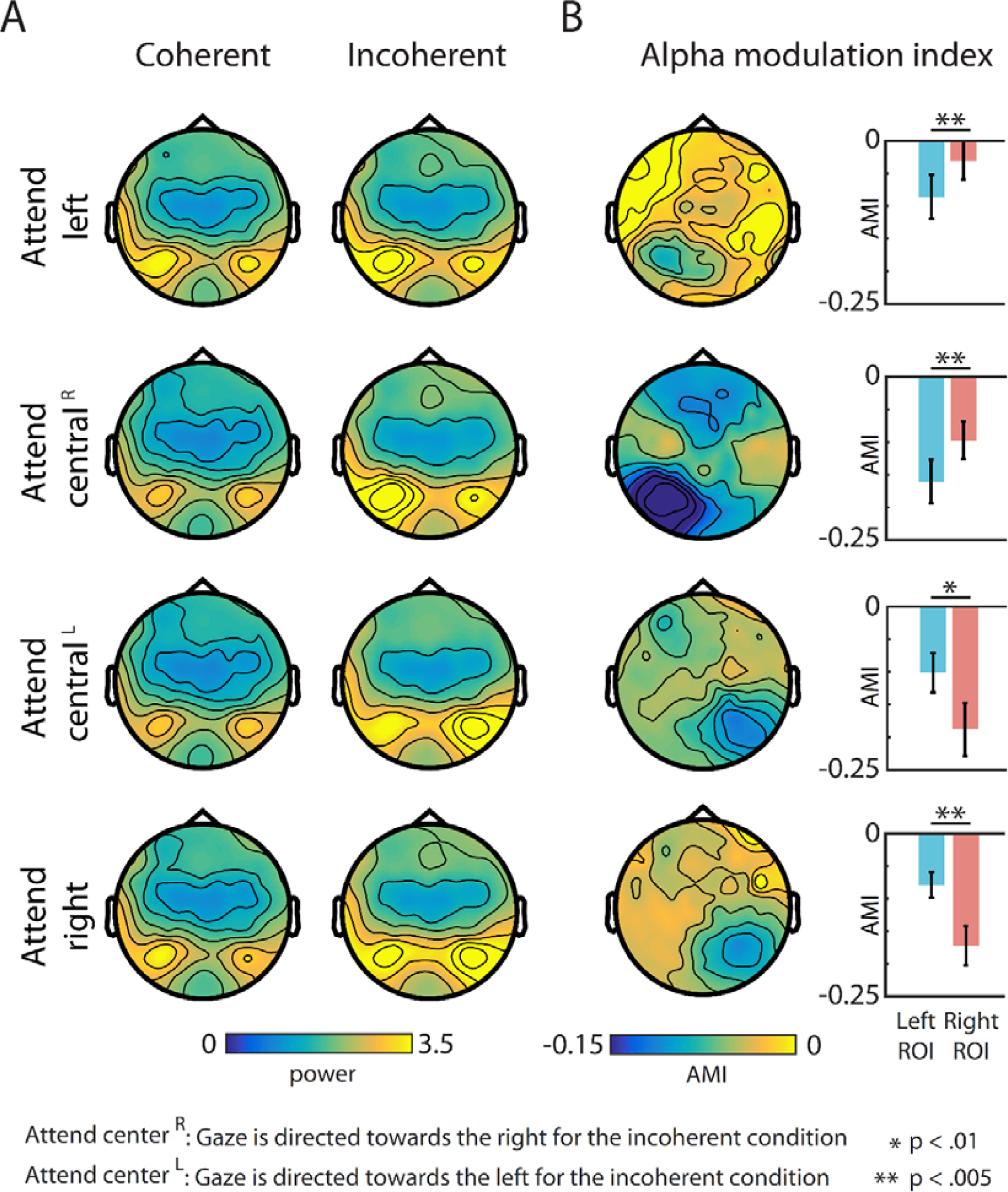
Topographies and statistical results for alpha-band power. **(A)** Topographies of alpha-band (8-14 Hz) power for the coherent (left column) and incoherent condition (right column). Occipital alpha-power is overall increased in the incoherent condition. Data are presented separately for conditions in which participants attended towards the left (top row), towards the center (middle two rows), and towards the right (bottom row). For attend central^R^ (upper middle row) gaze is directed towards the right in the incoherent condition. For attend central^L^ (lower middle row) gaze is directed towards the left in the incoherent condition. **(B)** Topographies (left column) and statistical results (right column) for the alpha-modulation 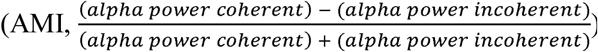. Data are presented separately for the attend-left, attend-central^R^, attend-central^L^, and attend-right. Occipital AMI was significantly lateralized for all attention locations, due to stronger alpha-power contralateral to the direction of gaze in the incoherent condition.

For statistical comparisons of AMI (Figure 7B), we calculated a 2-way ANOVA using the factors Attentional Location (*left* vs. *central^R^* vs. *central*^L^ vs. *right*) and ROI (*left* vs *right).* We did not find a main effect (both p > .059), but a significant 2-way interaction between the factors Attentional Location and ROI (F(3,54) = 24. 8, p < .001). Follow-up t-tests were calculated between AMI at the left and right ROI, for each Attention Location separately. For both the attend-left and attend-central^R^ condition, the t-tests yielded a significantly lower AMI in the left compared to the right ROI (t(18) = −3.7, p < .002 and (t(18) = −3.2, p < .005), resp.). For the attend-central^L^ and attend-right condition, the t-tests yielded a significantly lower AMI in the right compared to the left ROI (t(18) = 4.7, p < .000 and (t(18) = 5.2, p < .000), resp.). Further, AMI both ipsi-and contralateral to the ignored side differed significantly from zero (t(18) = −4.8, p < .017 and (t(18) = −6.7, p < .007), resp.). In summary, occipital alpha-power was significantly stronger in coherent compared to incoherent conditions, as shown by the AMI differing from zero. Alpha power was also significantly lateralized, shown by the difference in AMI between left and right ROIs.

### Theta-band results

Figure 8A depicts the results of the theta-band analysis. The topographical distribution of theta-band power was similar for the coherent and incoherent conditions, and strongest at fronto-central sites (Figure 8A, left- and middle topographical plots). Interestingly, the largest TMI values, as well as the statistically significant differences between the two conditions were found in more central and posterior areas (Figure 8A, right topographical plot). The first level statistical analysis using cluster-corrected pairwise t-tests between conditions revealed 36 channels exhibiting significant differences. In a second level analysis, we ran one-sample t-tests separately for each loudspeaker location, to test whether TMI differs from zero (Figure 8B). We found significant differences for the attend-central^R^, (t(18) = −2.6, p < .017) attend-central (t(18) = −2.3, p < .033) and attend-right (t(18) = −3.2, p < .005) condition, suggesting that theta power is larger in the incoherent compared to the coherent conditions. No significant difference was found for the attend-left condition (p > 0.11). Noticeably, we also found the numerically lowest (although statistically significant) alpha-band modulation for the attend-left condition. A potential, straightforward explanation for this finding is the layout of our experimental booth. Due to the spatial constraints of the experimental booth participants sat closer to the left compared to the right side of the room (0.7 and 1.7 m distance from the centre of the headrest to the left and right walls of the booth, respectively). While we took great care in the physical setup of the present experiment, it is possible that these asymmetries have caused acoustic differences between sounds presented from the left versus from the right which affected the observed theta band power. Finally, to investigate potential differences in TMI between attended loudspeaker locations, we computed a repeated measures ANOVA on mean TMI values, using the factor Attention Location (*left* vs. *centraf* vs. *centraf* vs. *right*). This test yielded no differences in TMI (p > .52).

**Figure 8:**
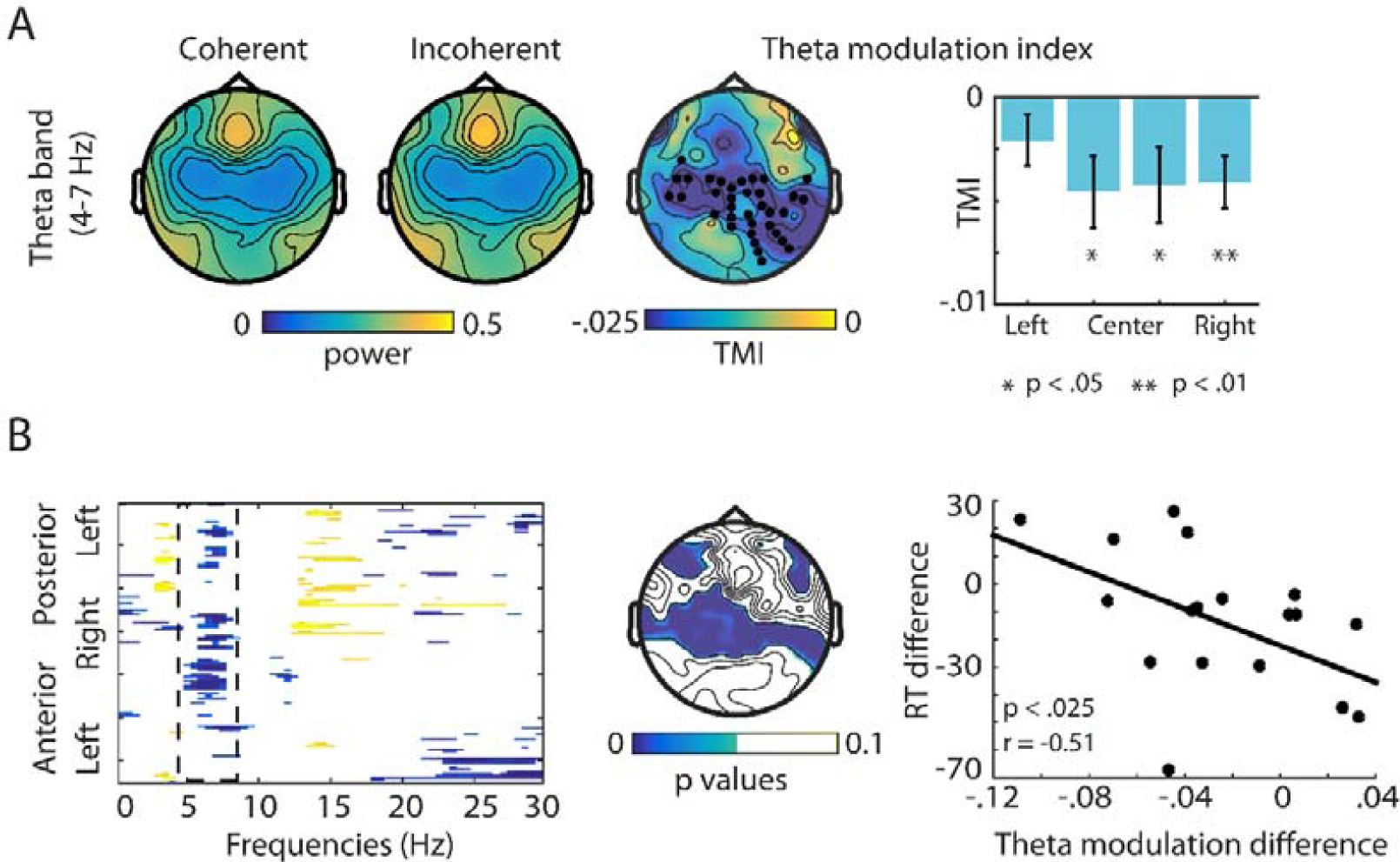
Theta-band results. **(A)** Results of theta-band (4-7 Hz) power analysis. Topographies are shown for the coherent (left), and incoherent condition (middle), as well as for the theta-modulation index 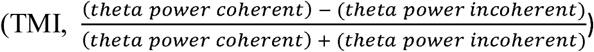. Black dots indicate the channels belonging to the observed significant cluster. The barplot shows the statistical results of testing TMI against zero, separately for the attend-left, attend-central^R^ attend-central^L^, and attend-right. **(B)** Exploratory correlation analysis between the effect of coherence between gaze and attention (coherent minus incoherent) in EEG power and response times. The left plot shows a map of the channel- and frequency wise point-by point correlation. The x-axis shows all frequencies from 1 to 30. Coded on the y-axis are all channels running from left-posterior (top) to left anterior (bottom), with each channel corresponding to one line. Only significant correlations are shown, with the correlation coefficient being coded by colour (blue = negative, yellow = positive). The middle plot illustrates the topography of all significant correlations for the theta band (4-7 Hz). For illustrative purposes, the left plot shows the correlation between the TMI and the response-times coherence effect for all significant channels.

### Correlation analysis results

To investigate a potential relationship between ongoing EEG activity and behavioural performance, we calculated an exploratory correlation of the effect of spatial coherence (i.e. coherent – incoherent condition) between spectral EEG power in all frequency bands from 2-30 Hz, and RTs. Figure 8B (left plot) displays a map of the significant (uncorrected) pointwise correlations for each channel and frequency. Of the different patches of significant correlations, a cluster in the theta band (4-7 Hz) had the largest number of significant channels (51) while being reasonably narrow banded in frequency. Since we also found significant power effects in the theta band, we selected this cluster for further investigation. Figure 8B, middle plot, shows the central scalp topography for this theta-band cluster (p-values masked for significance), which is similar to the topography of the theta power effect (Figure 8A). As a strictly illustratory measure, a scatterplot of the correlation within the significant channels of the theta band is depicted in Figure 8B, left plot.

## Discussion

In the present study we investigated the impact of task irrelevant visual gaze direction on auditory processing, and report three main findings. Gazing away from the locus of auditory attention leads to: (a) Increased RTs to attended sounds, indicating impeded auditory processing; (b) Increased occipital alpha-band power specifically contralateral to the direction of gaze, indicating a suppression of distracting input; (c) Overall increased central theta-band power, indicating extended recruitment of top-down cognitive control. Contrary to our hypothesis, we found no effect of gaze on auditory ERPs.

### Gaze affects behavioural responses to individual sounds

Independent of the attended spatial location, responses to targets were overall slower when participants gazed away from the attended location, compared to when they gazed toward it. Changes of gaze direction are typically used to align the fovea with the currently attended location in a visual scene. Thus, a potential straightforward explanation of our finding is that a misalignment of the locus of visual attention with the locus of auditory attention leads to impeded auditory processing. This is in line with previous studies reporting decreased accuracy in auditory spatial localization when gazing away from a sound source^9,43^ Maddox et al.^9^ suggest that the gaze-related changes in auditory spatial acuity are the result of a matching between visual and auditory spatial maps, mediated via crossmodal integration in midbrain structures such as the inferior and superior colliculus. In is conceivable that our present finding of increased RTs in the incoherent condition is also a consequence of gaze-related reduction in spatial acuity, which hampers the spatial separation of the three sound streams and delays target detection. Further, our results might also reflect that gaze affects the ability to ignore distracting sounds. Indeed, Spence and Driver^17^ have shown that distracting sounds are harder to ignore when they are visually fixated. Although their study was similar in design to our present one, Okita and Wei^29^ did not find gaze-related differences in RTs to auditory targets. A potential reason for this is that their task was overall easier (longer ISIs, only 2 loudspeaker locations, placed further apart), with performance close to ceiling level (94 % hit rate, 0.05 % FAs).

In addition to slower RTs, we also found less FAs in the incoherent compared to the coherent condition. This unexpected finding can be interpreted in the context of perceptual load theory^50,51^, which predicts that task irrelevant stimuli are easier to suppress during tasks which demand a high compared to a low perceptual load. The misalignment of gaze and auditory attention presumably constitutes a task with higher perceptual load than an alignment of gaze and attention, and protects from a spill-over of attentional resources from task-relevant to task irrelevant stimuli. Further, it is worth mentioning that our finding includes FAs occurring at any time during the experiment (i.e. responses following targets as well as standard sounds). When disregarding FAs to standards, and taking into account only responses following target sounds presented from the two currently unattended loudspeakers, FA’s did not differ between coherent and incoherent conditions (p > 0.05).

### Event-related potentials are unaffected by gaze

In line with the extensive literature on neural correlates of auditory attention^10–13^, we found a systematic increase of early ERP amplitudes to attended compared to unattended standard sounds, independent of concurrent gaze direction. Specifically, P1 amplitudes were overall larger for attended compared to unattended sounds across attentional locations, while N1 amplitudes were larger for attended sounds from the left and right but not the central location. The lack of a significant difference in the latter condition corresponds to the participants’ self-report that attending to the central location subjectively was the most difficult. This is also reflected in the significant behavioural FA main effect for location, presumably due to significantly higher FA rates for the attend centre compared to the attend-left and attend-right conditions (see Figure 4). Additionally, we found larger P2 amplitudes for unattended compared to attended sounds. Again, this is in line with previous studies on auditory attention, showing an increase of ERP amplitudes around 200 ms post-stimulus for actively ignored compared to attended sounds^52,53^. This increase is thought to reflect a process of distractor suppression, and has been shown to increase with training^54^.

As a main finding, and contrary to our hypothesis, we observed no effect of gaze on ERPs as well as no interaction between attention and gaze. This is in contrast to the results reported by Okita and Wei^29^, who, using a similar design, found a smaller negative difference wave (i.e. the difference between ERPs during attended minus unattended conditions) for not gazed-at versus gazed-at sounds. Importantly however, they suggest that this smaller attentional effect for the not gazed-at sounds reflects a decrease in the selectivity between relevant and irrelevant inputs, which fits well with the interpretation of our present behavioural and oscillatory data. Further, and contrary to both our and Okita and Wei’s^29^ findings, a recent study investigating the impact of gaze on somatosensory processing^55^ reported that gazing away from the location of an attended tactile stimulation led to an increase rather than a decrease in ERP amplitudes around the N140. The authors also report decreased behavioural performance in conditions where gaze was directed away from the somatosensory input, and suggest that the mechanisms of attention and gaze operate in parallel and are independently reflected in the concurrent neural processes. Together, these studies show that evidence on the effect of gaze on attention-related ERPs in non-visual domains is still scarce and inconsistent. However, it is generally thought that gazing away from the locus of attention causes distraction and hinders processing of the relevant input. Importantly, both Okita and Wei^29^ and Gherri and Forster^55^ employed easier tasks compared to our study. It is likely that the higher task difficulty in our experiment led to a more focused auditory selective attention, and concomitant suppression of sensory input (likely, both auditory and visual) from irrelevant spatial locations. This suppression of visual input, which was task irrelevant, might have reduced the impact of gaze on ERPs and is consistent with the strong increase in occipital alpha band activity we observed. Finally, our present null finding might be explained merely by a too low signal to noise ratio, and certainly does not rule out the general presence of a gaze mediated effect on auditory ERPs.

### Occipital alpha-oscillations reflect suppression of misaligned gaze information

While we observed no effect on evoked response, the analysis of ongoing oscillatory activity revealed large differences in brain state between ‘coherent’ and ‘incoherent’ conditions. Overall, in line with the behavioural data, it appears that significant energy was exerted to counteract the inconsistent gaze location (Figures 7 and 8). Occipital alpha-band power was consistently increased during the incoherent compared to the incoherent condition, as shown by a negative alpha-modulation index. This occipital alpha activity likely indicates the suppression of distracting, task irrelevant input, and has been demonstrated in numerous studies modulating attention toward auditory^18,21^, visual^22,45^, and multimodal stimuli^56,57^ Importantly, during the incoherent condition, the modulation of alpha power was stronger contralateral to the direction of gaze compared to ipsilateral. This was the case for all four attentional conditions (Figure 7B). This indicates that the increase in alpha power is not just an unspecific response to increased task demands or audio-visual mismatch, but a spatially distinct attention mechanism to supress information from one hemifield. Our results fit with previous findings of modulations in lateralized alpha band activity during covert spatial attention, both in the auditory^18^ and visual^45^ domain. Judging from the scalp topography it is difficult to assess whether this alpha modulation acts to suppress visual, auditory, or inputs from both modalities. However, the fact that our task involved dynamic auditory, but not visual stimulation from the unattended side might suggest that at least the lateralized, spatially selective suppression acts primarily on auditory inputs. Further, the difference in occipital alpha-band power between the coherent and incoherent conditions, and thus in the suppression of distracting input, might be an additional reason for the observed lower FA rate in the incoherent condition.

### Increased theta power reflecting top-down control during spatial incoherence

Apart from modulations in occipital alpha-band power, gazing away from the locus of auditory attention also increased central theta-band power (Figure 8A). Specifically, significant effects in theta-band modulation were found during the attend-central^L^ attend-central^R^ and attend-right conditions. Similar changes in central and fronto-central theta-band power have recently been linked to task requirements and cognitive control^24,26,58,59^ Clayton^25^ suggested that increasing levels of theta during prolonged attention reflect both increasing processing demands and resulting fatigue, as well as simultaneous compensatory upregulation of top-down control. In our present study, the observed concurrent increase in occipital alpha-band power might be a consequence of the theta-band mediated upregulation of top-down control during the more demanding incoherent condition. Two recent studies found similar combined effects in the alpha and theta-bands to what we report here. Ahveninen et al.^23^ compared top-down cue-directed auditory attention with novelty-based bottom-up attention during a dichotic listening task. The authors report increased ipsi-compared to contralateral alpha, as well as an increase in fronto-medial theta power during the cued spatial attention, indicative of increased underlying top-down processes. Combined effects of posterior alpha and anterior theta modulation in response to processing demands were also found by van Noordt et al.^58^ during a cued saccade versus antisaccade task. Finally, our exploratory correlation analysis revealed a negative relationship between the benefits of gazing toward the attended location in RTs and theta-band power. That is, a slowing down of RTs during the incoherent condition was associated with a stronger increase in central theta-band power. This finding corresponds well with the theta-band literature cited above, and suggests a link between theta-band power and task demands. Participants who were particularly affected by the spatial incongruence between gaze and auditory attention increased top-down cognitive control acted as a compensatory mechanism. Alternatively, participants who used equal levels of cognitive control in both coherent and incoherent conditions consequently produced faster RTs in the less demanding coherent condition. This is in line with previous studies reporting correlations between theta-band power and error rates during sustained attention^60^ as well as between theta-band power and subjective mental effort^26^. Despite their correspondence with our power results as well as with the literature, it is, important to stress that the present correlation analyses was exploratory in nature, and the reported effects are based on mass statistical tests analysis, uncorrected for multiple comparisons.

### Conclusion

Here, we provide evidence that gazing away from the location of auditory attention leads to impeded processing of sounds. While we did not find earlier reported effects of gaze direction on ERPs to individual sounds, to our knowledge this is the first report showing effects of gaze direction on behavioural auditory target detection and concurrent oscillatory activity in the alpha and theta-band range. It is likely that a spatial mismatch between visual gaze and auditory attention leads to increased task demands, as reflected in slowed RTs during the incoherent condition. Stronger occipital alpha-band power contralateral to the direction of gaze is indicative of a spatially selective suppression of task irrelevant information. Further, increased central theta-band activity likely reflects enhanced cognitive control mechanisms, which correlate with behavioural effects and potentially mediate the observed increase in alpha-power. It is possible that both alpha- and theta-band mediated compensatory processes were largely successful at eliminating the adverse effects of inconsistent gaze, thus explaining the lack of effects of gaze onto ERPs to individual sounds. While these compensatory mechanisms might work well in our cohort of young participants, inconsistent gaze might have more severe consequences in older and/ or hearing impaired listeners. Finally, our results highlight the potential impact of task irrelevant low-level visual input on auditory processing, and demonstrate the importance of proper visual fixation control in studies on auditory attention.

## Acknowledgements

We are grateful to the SR research support team for help with the eyetracking setup. This research was supported by a EC Horizon 2020 grant (MC).

## Author contributions

UP and MC designed the experiment. UP collected and analysed the data. UP and MC wrote the manuscript.

